# Development of a cricket paralysis virus-based system for inducing RNA interference-mediated gene silencing in *Diaphorina citri*

**DOI:** 10.1101/2020.11.15.383588

**Authors:** Emilyn E. Matsumura, Jared C. Nigg, Elizabeth M. Henry, Bryce W. Falk

## Abstract

*Diaphorina citri*, the Asian citrus psyllid, is the insect vector of the phloem-limited bacterium ‘*Candidatus* Liberibacter asiaticus’, which causes the most devastating citrus disease worldwide: Huanglongbing (HLB). An efficient cure for HLB is still not available and the management of the disease is restricted to the use of pesticides, antibiotics and eradication of infected plants. Plant- and insect-infecting viruses have attracted increasing attention for their potential to manipulate traits in insects, especially insect vectors of plant pathogens. However, so far there are no insect virus-based vectors available for use in *D. citri*. Cricket paralysis virus (CrPV) is a well-studied insect-infecting dicistrovirus with a wide host range and has been used as a model in previous translational studies. In this work, we demonstrate for the first time that CrPV is infectious and pathogenic to *D. citri.* We show that specific amino acid mutations in the CrPV primary cleavage DvExNPGP motif resulted in a viral mutant that was attenuated compared to wild-type CrPV during infection of either *Drosophila* cells line or adult *D. citri* insects. This attenuated CrPV mutant was then used as the backbone for engineering a recombinant CrPV-based vector to specifically alter *D. citri* gene expression via the RNA interference (RNAi) pathway, a technology called Virus Induced Gene Silencing (VIGS). As proof-of-concept, we engineered recombinant CrPV-based vectors carrying nucleotide sequences derived from a previously reported *D. citri* target gene: the inhibitor of apoptosis gene (IA). RT-qPCR analysis of insects either microinjected or fed with the recombinant CrPV mutants showed decreased IA gene expression as soon as viral replication was detected, indicating that the engineered CrPV-based VIGS system enables functional gene silencing in *D. citri*. This novel insect virus-based tool is easily amenable to genomic modification and represents a technical advance for understanding interactions between insect virus-based VIGS systems and *D. citri.*

## Introduction

*Diaphorina citri* Kuwayama (Hemiptera: Liviidae), also known as the Asian citrus psyllid, is the insect vector of the phloem-limited bacterium ‘*Candidatus* Liberibacter asiaticus’ (*C*Las). *C*Las is the causal agent of the most devastating citrus disease worldwide: Huanglongbing (HLB) or Citrus Greening [1, 2]. In the absence of an effective cure for HLB, disease management currently relies on insecticides, antibiotics and eradication of infected citrus plants [3–5]. However, these current HLB management practices also create environmental and public health concerns. Insecticides, for example, are not specific to the psyllid vector and might be harmful to beneficial insects [6]. Likewise, previous works have demonstrated that the application of antibiotics in citrus plants also alters the composition and reduces the diversity of the bacterial community present in citrus leaves [2, 7]. In addition, insecticide resistance has been previously reported for *D. citri* populations in Brazil and in the United States [3, 8]. Thus, these concerns highlight the urgent need for alternative psyllid management practices that facilitate reduced use of insecticides and antibiotics and/or a rotation between chemical application and other management methods with different modes of action [3].

RNA interference (RNAi)-based technology has shown to be an effective approach in controlling agriculture pests [9]. The RNAi pathway in insects is well documented [10–13], and most of the RNAi-based strategies use double-stranded RNA (dsRNA) to trigger RNAi responses in the target insects [14–21]. Several previous works have already demonstrated successful use of RNAi technology to specifically alter *D. citri* gene expression [4, 22–24], mostly by oral delivery of *in vitro* synthesized dsRNAs specific to *D. citri* genes [4, 23, 25–29]. However, the use of dsRNA as an inducer of RNAi in insects has shown some limitations due to its low stability in the natural environment and for eliciting an RNAi-response that is constrained to the insect gut [30]. Additionally, the choice of the strategy employed for delivering dsRNA into the target insect has also been a challenge [13].

A promising alternative is the use of insect viruses as vectors to deliver dsRNAs to target insects, a technology called Virus Induced Gene Silencing (VIGS) [31–34]. Upon virus infection, RNAi-mediated antiviral defense is induced in the insects, which targets both viral RNA and any transgenic RNA inserted within the virus genome [31–34]. In attempting to explore insect viruses as potential VIGS vectors in *D. citri*, several *D. citri*-specific viruses were discovered through next generation sequencing [35]. However, established infectious clones for any of these previously described *D. citri*-specific virus are still unavailable.

Cricket paralysis virus (CrPV) is a well-studied insect virus that has a wide host range [36–41] and established infectious clones are available [42]. CrPV belongs to the family *Dicistroviridae* and has a positive single-stranded RNA genome containing two open reading frames (ORFs), both translated from internal ribosome entry sites (IRESs) [43–45]. In this work, we show that CrPV also infects *D. citri.* We used a previously constructed CrPV infectious clone (a gift from Dr. Shou-Wei Ding) to develop a recombinant CrPV-based VIGS system carrying transgene sequences that specifically target *D. citri* through RNAi-mediated gene silencing. This is a valuable insect virus-based tool that can be used for several applications, such as (1) to study the interactions between an insect virus-based VIGS systems and *D. citri*, (2) to screen for *D. citri* or even *C*Las target genes, and (3) to establish an efficient delivery method for the VIGS system to other target insects. Additionally, the knowledge obtained from studies using the CrPV-based VIGS system in *D. citri* could be translated to naturally-infecting viruses of *D. citri* (Nouri et al., 2016) in attempts to develop specific, and perhaps field-applicable virus-based tools to complement HLB management.

## Results

### CrPV is infectious and pathogenic to D. citri

To assess the infectivity of CrPV in *D. citri*, CrPV virions were produced in *Drosophila melanogaster* Schneider line 2 (S2) cells after transfection with *in vitro*-transcribed RNAs (Fig S1A) derived from a previously constructed CrPV infectious clone (kindly provided by Dr. Shou-Wei Ding). The accumulation of CrPV RNA in transfected S2 cells due to viral replication was demonstrated by northern blot analysis (Figure S1B); and purified CrPV virions were examined by sodium dodecyl sulfate polyacrylamide gel electrophoresis (SDS-PAGE; Figure S1C) and transmission electron microscopy (TEM, Figure S1D). The infectivity of purified CrPV virions was further confirmed by inoculating S2 cells with purified virions and checking for CrPV-induced cytopathic symptoms (not shown).

The purified CrPV virions were used for microinjection of adult *D. citri* insects (~ 20 days-old). Insects were intrathoracically microinjected with either ~ 300 nL of a 10^3^ TCID_50_ units/μL virus suspension in 10 mM tris buffer (pH 7.4) or virus-free buffer as control. Microinjected insects were maintained in a single citrus leaf system and monitored daily for mortality or symptom development (Figure 1A). By the third day post injection (dpi), either paralysis symptoms or mortality were observed in insects microinjected with CrPV virions (Figures 1B and 1C). By day 6, approximately 90% of the insects injected with virus suspension were dead, while insects injected with virus-free buffer had an approximate mortality rate of 35% (Figure 1B). The accumulation of CrPV RNA in injected insects was monitored over time by RT-qPCR using primers to detect the CrPV coat protein (CP) gene. Total RNA from three replicate pools of three insects per pool was analyzed at time 0 (just after injection) and at 3 and 6 dpi. Viral genome copy number over time showed a gradual increase in CrPV-injected insects, being approximately 5 log10 units higher at 6 dpi compared to time 0 (Figure 1D). The increased RNA accumulation was also demonstrated by northern blot analysis (Figure 1E). These results confirm that CrPV is infectious and severely pathogenic to *D. citri* when delivered by intrathoracic microinjection.

**Figure 1:**
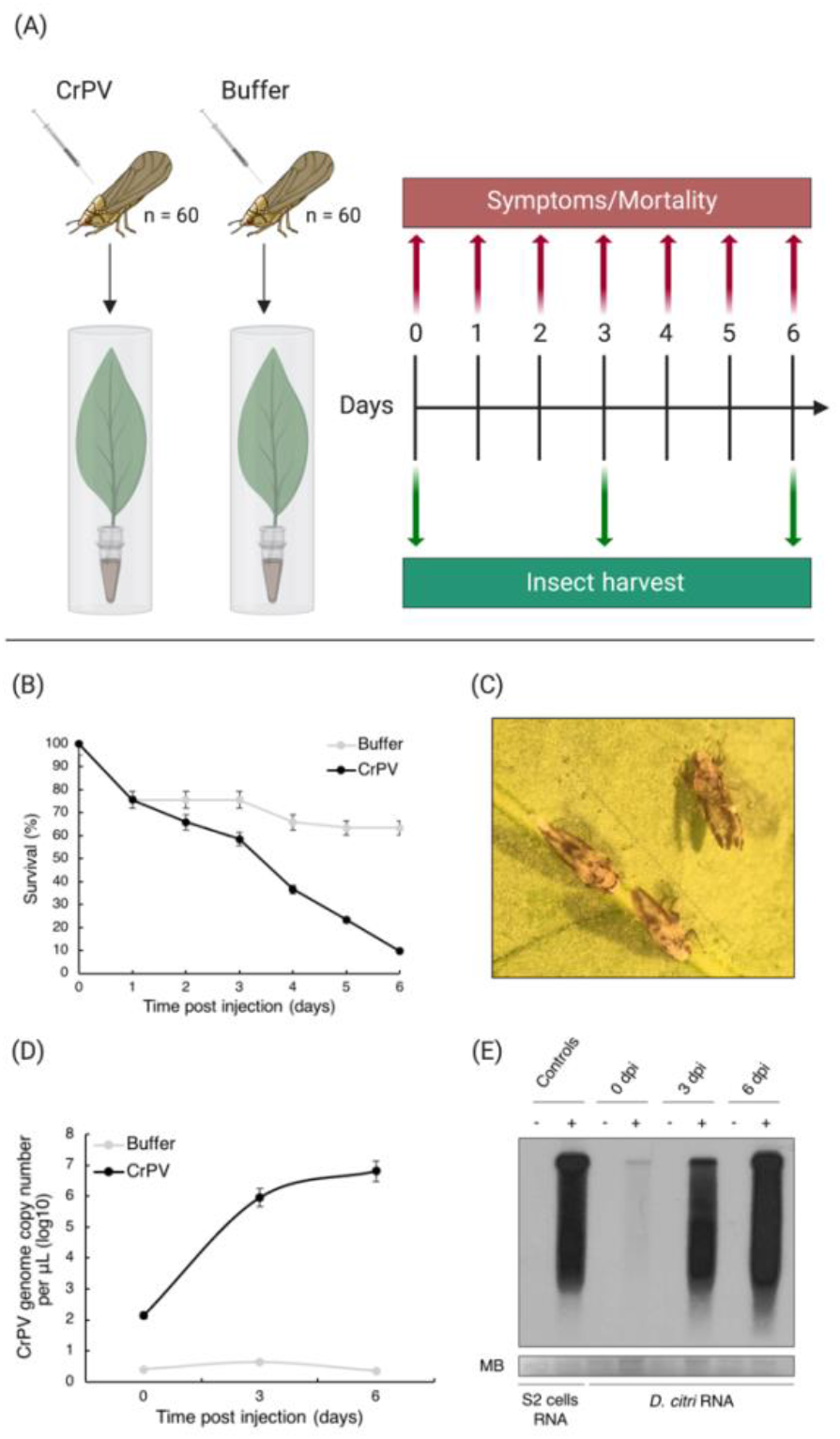
Infectivity of Cricket paralysis virus (CrPV) in *Diaphorina citri*. (A) Insects were intrathoracically microinjected with either purified CrPV virions or buffer solution (as control). Microinjected insects were maintained in a single leaf system for six days and were monitored daily for mortality and symptoms; n, number of insects used. (B) Survival curve of adult insects injected with CrPV. Survival rates were monitored at three time points post injection. Three biological replicates of n = 20 insects each per condition were analyzed. Error bars indicate standard error of the mean. (C) Cluster of flightless insects (a known paralysis symptom) observed in CrPV-injected *D. citri* under light microscope. Viral RNA accumulation was checked over a time-course by RT-qPCR (D) and confirmed by northern blot analysis on the total RNA from the injected insects (E). RNA from CrPV- or mock-infected S2 cells was used as a control. In (D), viral RNA accumulation is expressed as log10 genome copy number per μL (10 ng of RNA), and each time point represents data from three replicates of three pooled insects, and error bars indicate standard error of the mean. (−) Mock, buffer-injected insects; (+) CrPV-injected insects; dpi, days post injection; MB, methylene blue staining.

### Mutations in the primary cleavage motif of CrPV resulted in attenuation

CrPV infection in *D. citri* caused nearly 100% mortality by six days post infection when delivered by intrathoracic microinjection, thus complicating the use of CrPV as a vector for VIGS. The CrPV 1A protein is a potent suppressor of RNAi silencing [46], and likely a pathogenicity factor, therefore we sought to construct an attenuated variant of CrPV by mutating the conserved DvExNPGP motif, which is responsible for the co-translational cleavage of the 1A protein during translation of the ORF1-encoded polyprotein [46]. The wild-type (WT)-CrPV infectious clone was used as a template for the introduction of site directed mutations within the conserved DvExNPGP motif, which corresponds to nucleotides (nts) 1186-1209 in the CrPV genome (Figure 2A). In this experiment, we examined the effects of both single and double amino acid (aa) mutations. Figure 2B shows all the mutants constructed for this analysis.

**Figure 2:**
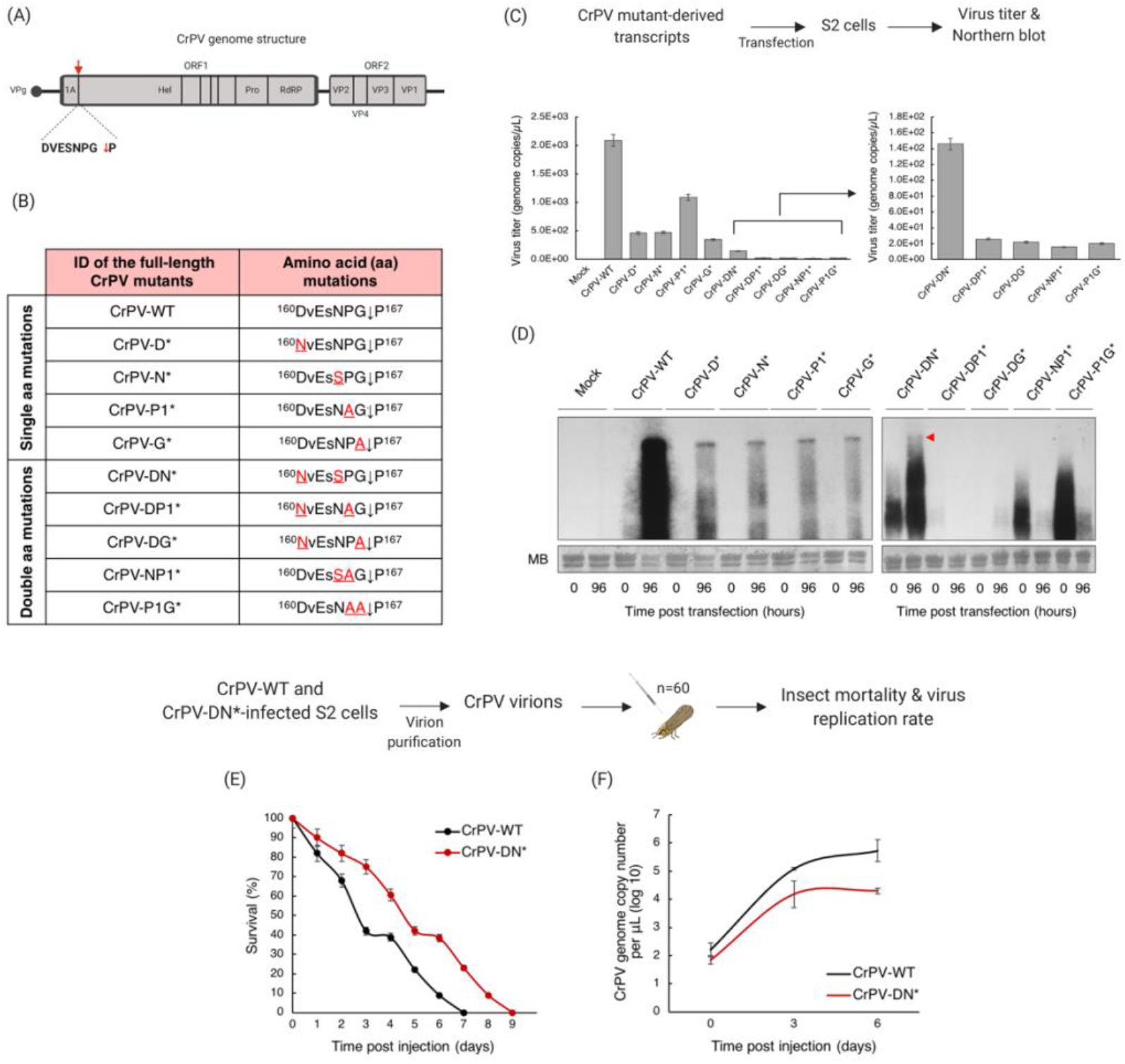
Construction of an attenuated variant of Cricket paralysis virus (CrPV) by mutational analysis on the CrPV DvExNPGP motif: a signal motif for cleavage of the 1A silencing suppressor protein. (A) Schematic representation of the CrPV genome organization showing the location of the DvExNPGP motif as well as the cleavage site (red arrow). (B) List of single and double amino acid (aa) mutations introduced into CrPV DvExNPGP motif; mutated aa’s are highlighted in red. CrPV mutant-derived transcripts were transfected into S2 cells and viral RNA accumulation in the supernatant was evaluated 96 hours post transfection by both RT-qPCR (C) and northern blot analysis (D). In (C), viral RNA accumulation is expressed as log10 genome copy number per μL (10 ng of RNA), and error bars represent standard error of the mean of four biological replicates. The infectivity of the CrPV-DN* mutant (B), selected as an attenuated variant of CrPV, was evaluated in *Diaphorina citri* by microinjecting CrPV-DN*-derived virions into adult insects. Microinjected insects were monitored daily for mortality (E). Three biological replicates of n = 20 insects each per condition were analyzed. Error bars indicate standard error of the mean. Viral replication rates were checked over a time-course by RT-qPCR (F). Each time point represents data from three replicates of three pooled insects, and error bars indicate standard error of the mean. For comparison, this experiment was simultaneously performed with wild-type (WT) CrPV. MB, methylene blue staining; n, number of insects used.

The viability of CrPV carrying these mutations was first evaluated in *Drosophila* S2 cells by transfection of *in vitro*-transcribed RNAs derived from the mutated clones. Viral replication (viral RNA accumulation) and induction of cytopathic effects was monitored during infection with each mutant in S2 cells. Although the single aa mutants CrPV-D*, CrPV-N*, CrPV-P1* and CrPV-G* (Figure 2B) were still able to replicate in the transfected cells, accumulation of viral RNA in the supernatant at 96 hours post transfection (hpt) was reduced by 78, 77, 48 and 83%, respectively, relative to WT-CrPV control (Figures 2C and 2D). These results demonstrate that the respective aa mutations negatively impacted CrPV replication. However, the cytopathic effects induced by infection with these mutants in S2 cells were very similar, in both severity and time of appearance, to the effects induced by the infection of WT-CrPV (data not shown).

We then evaluated the viability of the CrPV mutants harboring double aa mutations in the DvExNPGP motif. Five independent double aa CrPV mutants were constructed (Figure 2B). Only one double aa mutant (mutant CrPV-DN*) replicated in S2 cells. The mutations introduced in the CrPV genome to generate mutants CrPV-DP1*, CrPV-DG*, CrPV-NP1* and CrPV-P1G* (Figure 2B) resulted in non-viable viral variants (Figures 2C and 2D). Accumulation of viral RNA in the supernatant of S2 cells transfected with CrPV-DN* RNA was reduced by 93% at 96 hpt compared to cells transfected with WT-CrPV RNA (Figures 2C and 2D). Furthermore, the appearance of cytopathic effects in cells transfected with *in vitro*-transcribed RNAs derived from the CrPV-DN* mutant was delayed by 2 days, compared to cells transfected with WT-CrPV-derived transcripts (data not shown).

Due to the delay in cytopathic effects in S2 cells, the CrPV-DN* mutant was selected for testing in *D. citri*. Approximately 300 nL of a 10^3^ TCID_50_ units/μL of either CrPV-DN* or WT-CrPV purified virions were intrathoracically microinjected into adult insects, which were then kept in a single leaf system and monitored daily for mortality. The CrPV-DN* and WT-CrPV replication rates in the injected insects were also evaluated by a time course RT-qPCR assay. Although the CrPV-DN* mutant still showed the ability to replicate in and kill the injected *D. citri* insects, a delay in insect mortality was observed when compared to the mortality rate of WT-CrPV-injected insects (Figure 2E). It is likely that this observed delay is due to the lower replication rate of the CrPV-DN* mutant in these infected insects, as shown by the time course RT-qPCR analysis (Figure 2F). These results indicate that the mutant CrPV-DN* is attenuated compared to WT-CrPV in both S2 cells and *D. citri*. Thus, mutant CrPV-DN* was selected as the virus-based vector for use in further VIGS experiments in *D. citri*.

### A recombinant CrPV-based VIGS system was established to specifically target D. citri

To evaluate CrPV as a viral vector for VIGS experiments, the mutant CrPV-DN* infectious clone was modified to harbor inserts derived from the selected target gene. Sequences were inserted downstream of the CrPV DvExNPGP motif (modified to NvExSPGP in the CrPV-DN* mutant), and a duplicated DvExNPGP motif was inserted downstream of the insertion site to also release the ORF1-encoded polyprotein from the extra aa’s translated from the insert (Figure 3A). To determine the maximum insert length CrPV can tolerate within its genome, different lengths of green fluorescent protein (GFP) gene-derived sequences (Figure 3B) were individually inserted into the CrPV-based vector, and the stability of recombinant CrPV was evaluated in S2 cells. CrPV virions carrying GFP-derived sequences of 57, 102 and 150 nt in length were retained during infection in S2 cells (Figure 3C) and the recombinant viruses were assembled into virions indistinguishable from WT-CrPV virions (Figure 3D). However, a recombinant virus carrying an insertion of a 201 nt GFP-derived sequence was not viable. Thus, an insert length of 150 nt was selected as the optimum insert length for the CrPV-based VIGS system.

**Figure 3:**
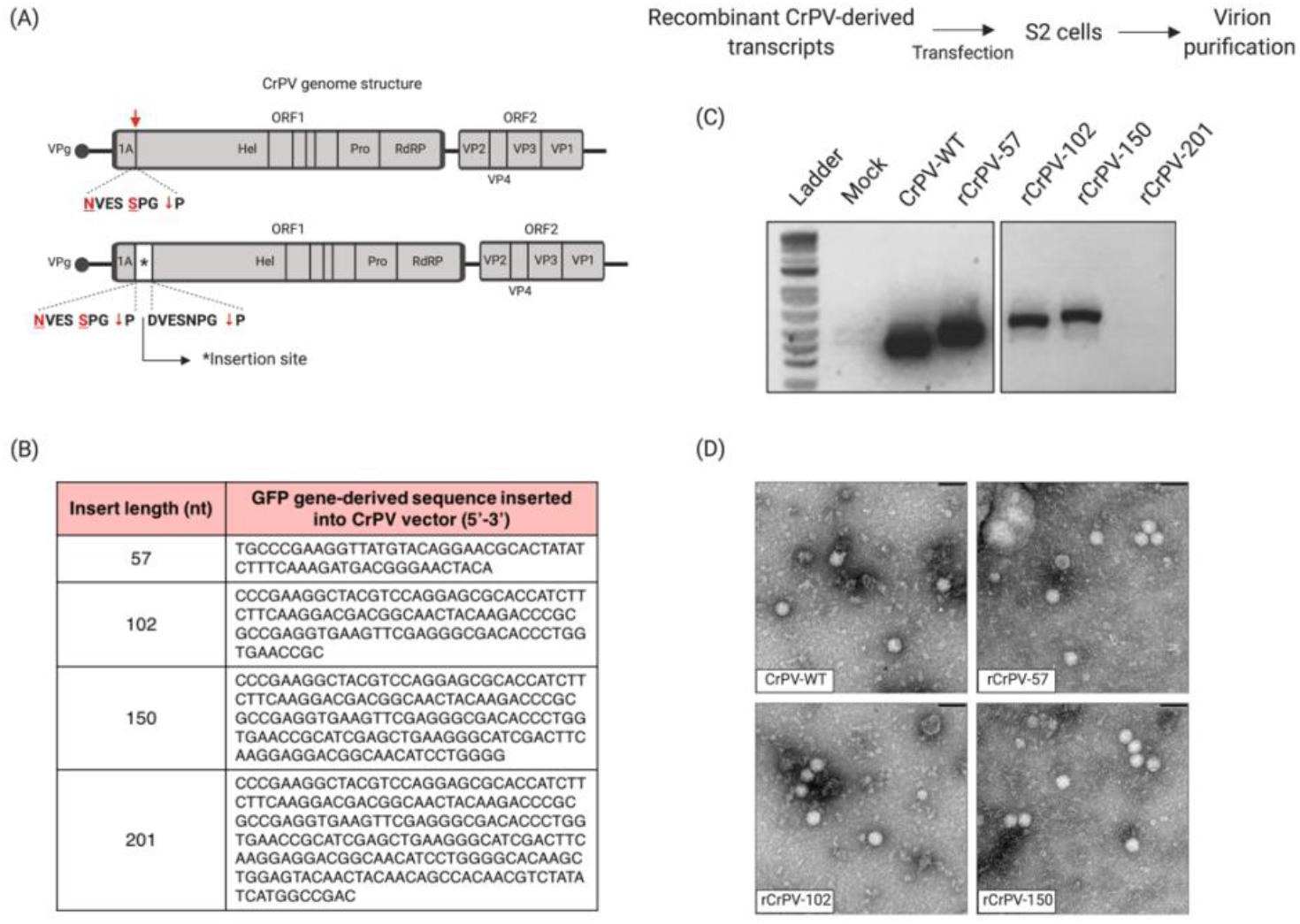
Construction and stability analysis of the recombinant clones of Cricket paralysis virus (CrPV). (A) Schematic representation of the genome structures of CrPV-DN* mutant (from this work, top) and its derivative recombinant CrPV (rCrPV, bottom) showing the insertion site for cloning the desired inserts. (B) List of the green fluorescent protein (GFP) gene-derived sequences (shown as DNA sequences) individually inserted into the CrPV insertion site to generate the rCrPVs; the respective nucleotide (nt) lengths of each sequence are shown. Recombinant CrPV virions were recovered from *Drosophila* S2 cells transfected with the rCrPV-derived transcripts. Stability of the rCrPVs was evaluated by checking insert retention in the virion RNA, by RT-PCR (C); and by transmission electron microscopy of the rCrPV virions (D). Scale bars in (D) are 60 nm.

To establish proof-of-concept for the CrPV-based VIGS system in *D. citri*, a previously reported *D. citri* target, the inhibitor of apoptosis gene (IA) [4], was chosen as the target for this experiment. Since we demonstrated that 150 nts is the maximum insert length that can be carried by the CrPV-based vector, we first performed a dsRNA-mediated gene silencing experiment to determine whether short IA gene-derived sequences can still trigger RNAi responses in *D. citri* and, consequently, decrease the expression of the respective gene. The sequences for dsRNA synthesis were selected based on the *D. citri* IA gene sequence available in GenBank (Accession number: XM_008469919; Figure 4A), and the same sequences were used for engineering the recombinant CrPVs (Figure 5B). DsRNAs (150 bp) were *in vitro* synthesized considering both forward (IA-F) and reverse (IA-R) orientations of the IA gene. A 150 nt GFP gene-derived dsRNA was also synthesized to be used as a control. The size and integrity of the *in vitro* synthesized dsRNAs was analyzed on 1% agarose gel electrophoresis (Figure 4B) and delivered to *D. citri* insects either by microinjection or oral feeding, as described in *Methods*. Relative levels of IA transcripts were assessed by RT-qPCR. Insects injected with either dsRNA-IA-F or dsRNA-IA-R showed approximately 50% decrease in IA gene expression levels (p < 0.005) at 17 hours post injection (hpi), compared to control insects (injected with dsRNA-GFP; Figure 4C). At this same time point (17 hpi), slightly higher mortality rates were observed in insects injected with dsRNA-IA-F (approximately 5% higher compared to control) and with dsRNA-IA-R (approximately 10% higher compared to control), however, these differences were not statistically significant and no mortality was observed from this time point until the last monitored time point, at 160 hpi (Figure S2). On the other hand, relative expression of the IA gene in insects fed for 5 days on artificial diets containing either dsRNA-IA-F or dsRNA-IA-R showed approximate decreases of only 25% (p < 0.0005) and 35% (p < 0.05), respectively, compared to the control (Figure 4D), and no differences in mortality rate were observed between dsRNA-IA-F/R- and dsRNA-GFP-treated insects (Figure S2). These results indicate that although the decreased expression levels obtained either by injecting (50%) or feeding (25-35%) dsRNAs to *D. citri* did not cause significant insect mortality in tested insects, the 150 nt sequences derived from the *D. citri* IA gene and selected for insertion into the recombinant CrPV clones are functional in silencing IA gene expression.

**Figure 4:**
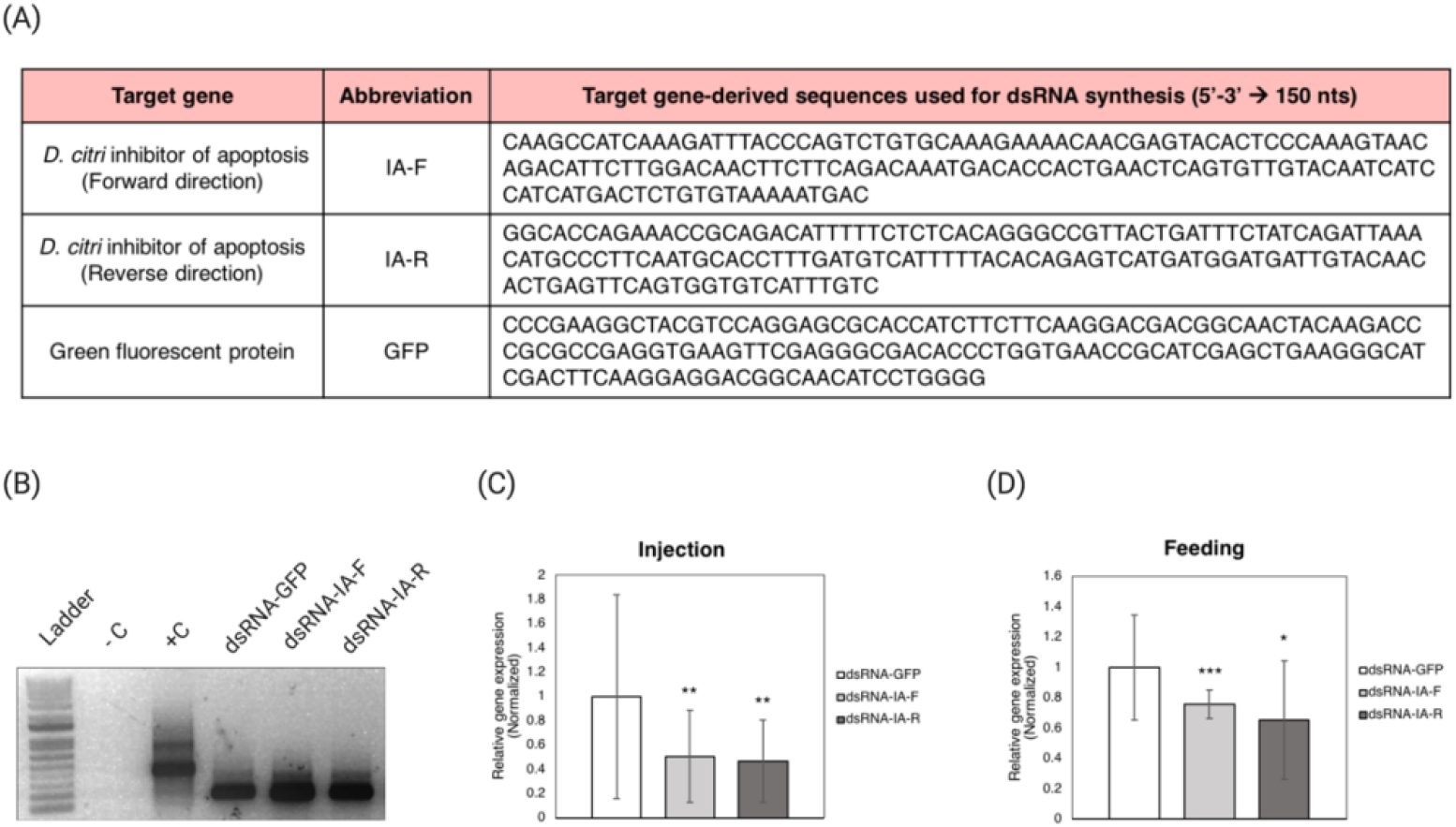
Gene silencing validation using *in vitro* synthesized dsRNAs (150 bp) corresponding to the *Diaphorina citri* inhibitor of apoptosis (IA) gene. (A) IA gene and green fluorescent protein (GFP) gene sequences (shown as DNA) used for the dsRNAs synthesis. (B) Agarose gel electrophoresis analysis of the *in vitro* synthesized dsRNAs. Ladder is 1 Kb plus DNA ladder; -C is a negative control; +C is a positive control (from MEGAscript RNAi kit). (C) IA gene expression analysis of *D. citri* insects injected with 200 ng of IA gene-derived dsRNAs; this analysis was performed 17 hours post injection by RT-qPCR. (D) IA gene expression analysis of *D. citri* insects fed on artificial diet containing 300 ng/μL of IA gene-derived dsRNAs; this analysis was performed after 5 days of feeding by RT-qPCR. Insects injected or fed with GFP gene-derived dsRNAs were used as controls and set as 1. IA gene-derived RNA levels were normalized against the *D. citri* actin gene. Bars represent the average IA gene RNA level of ten individual insects. Error bars indicate the standard error of the mean. * = p < 0.05, ** = p < 0.005, *** = p < 0.0005 (two-tailed t-test).

**Figure 5:**
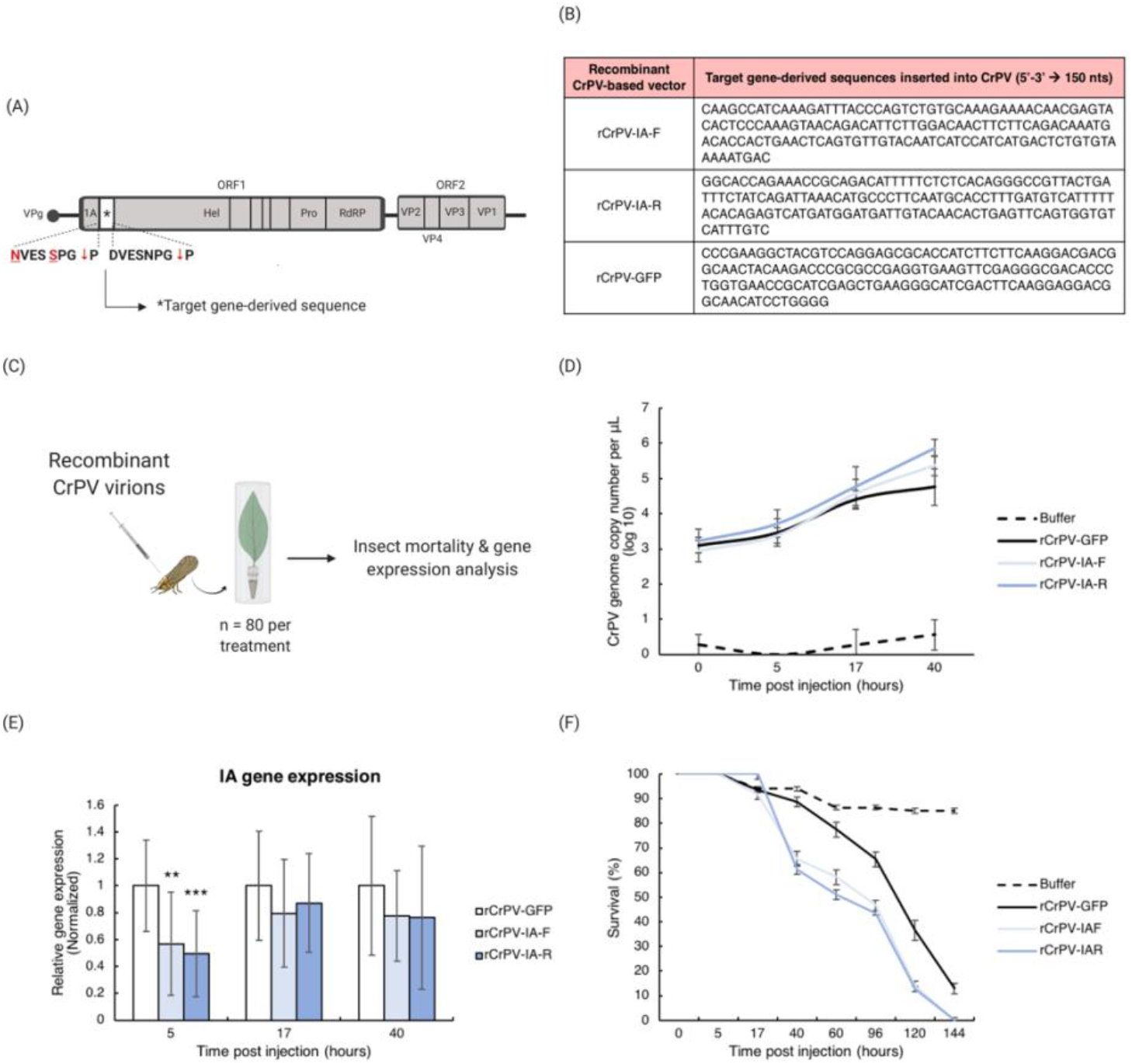
Virus-induced gene silencing (VIGS) using the recombinant Cricket paralysis virus (rCrPV) system for inducing RNA interference in *Diaphorina citri* by microinjection delivery. (A) Schematic representation of the rCrPV genome showing the insertion site of the target gene-derived sequences. (B) *D. citri* inhibitor of apoptosis (IA) gene- and green fluorescent protein (GFP) gene-derived sequences (shown as DNA) that were individually inserted into the rCrPV-based vector. (C) rCrPV virions (approximately 300 nL of a 10^3^ TCID_50_ units/μL) were intrathoracically injected into adult insects (n = 80), which were transferred to a single citrus leaf system until further analysis. (D) Viral RNA accumulation in rCrPV-injected insects was checked at three time points post injection by RT-qPCR; injected insects from time 0 (just after injection) were used to calculate the input of virions. Buffer-only injected insects were used as controls. Viral RNA accumulation is expressed as log10 genome copy number per μL (10 ng of RNA). Each time point represents the average from ten individual insects; error bars indicate the standard error of the mean. (E) IA gene expression analysis of insects injected with rCrPVs. This analysis was performed at three time points post injection by RT-qPCR. Insects injected with rCrPV-GFP were used as controls and set as 1. IA gene-derived RNA levels were normalized against the *D. citri* actin gene. Bars represent the average of IA gene RNA level from ten individual insects. Error bars indicate the standard error of the mean. ** = p < 0.005, *** = p < 0.0005 (two-tailed t-test). (F) Survival curve of adult insects injected with rCrPVs. Survival rates were monitored at eight time points post injection. Three biological replicates of n = 20 insects each per condition were analyzed. Error bars indicate standard error of the mean.

The IA gene-derived sequences (IA-F and IA-R) used for dsRNA synthesis (Figure 4A) were PCR amplified and cloned into the insertion site of the CrPV-based vector, flanked by the two cleavage motifs (5’ NvExSPGP and DvExNPGP 3’) as previously described (Figure 5A), generating the clones rCrPV-IA-F and rCrPV-IA-R (Figure 5B). The virions derived from the new recombinant CrPVs were recovered in S2 cells and verified by SDS-PAGE (Figure S3A). Insert retention in the virions was confirmed by RT-PCR (Figure S3B) and Sanger sequencing. The purified rCrPV virions were then used in VIGS experiments to specifically target the *D. citri* IA gene, and the rCrPV carrying a GFP gene-derived sequence (150 nt) was used as control.

### Recombinant CrPVs induced target gene downregulation in D. citri as soon as viral replication was detected

Approximately 300 nL of a 10^4^ TCID_50_ units/μL solution of the recombinant CrPV-derived virions (rCrPV-IA-F, rCrPV-IA-R and rCrPV-GFP) were delivered to *D. citri* insects (~20 days-old; n = 80 for each treatment) by intrathoracic microinjection. Injected insects were maintained in a single citrus leaf system and collected at several time points for gene expression analysis (by RT-qPCR) and monitored for mortality (Figure 5C). Viral RNA as well as IA gene mRNA levels in microinjected insects were checked at 5, 17 and 40 hpi (Figures 5D and 5E), while insect mortality was monitored until 144 hpi (when rCrPV-injected insects achieved 100% mortality; Figure 5F). Analysis of viral RNA levels showed gradual increases in the genome copy numbers of the rCrPVs over the time course, an indication of viral replication (Figure 5D). The replication rates for rCrPV-IA-F, rCrPV-IA-R and rCrPV-GFP were similar until 17 hpi, however, at 40 hpi, insects injected with the rCrPV-IA-F and rCrPV-IA-R virions showed slightly higher viral genome copy numbers (approximately 0.6 and 1.1 log10 units higher, respectively) compared to rCrPV-GFP-injected insects (Figure 5D), even though equivalent amounts of rCrPV-IA-F, rCrPV-IA-R and rCrPV-GFP virions were delivered to the insects (time 0 in figure 5D). If high CrPV titer is positively correlated with *D. citri* mortality, these results align with the higher insect mortality rate observed at 40 hpi for insects injected with rCrPV-IA-F and rCrPV-IA-R virions (34.4% and 38.7%, respectively), compared with the mortality rate of insects injected with rCrPV-GFP virions (11.4%; Figure 5F). Significant decreases (p < 0.05) in the IA gene expression level (approximately 50% in both rCrPV-IA-F- and rCrPV-IA-R-injected insects) were obtained only at 5 hpi (Figure 5E). At 17 and 40 hpi, although IA gene expression showed a decrease of 20% in both rCrPV-IA-F- and rCrPV-IA-R-injected insects, compared with rCrPV-GFP-injected insects, the difference was not statistically significant. There were also no significant differences in the IA gene expression levels between the rCrPV-IA-F- and rCrPV-IA-R-treated insects, suggesting that the viral vectors carrying the target gene-derived sequences in either forward or reverse orientation have equivalent efficacy in silencing the respective gene. These results indicate that the rCrPV-based VIGS tool, when delivered by microinjections, is functional in inducing IA gene silencing in *D. citri* at early time points post infection, but as viral replication increases in the insects, significant differences in IA gene-derived mRNA levels in tested insects are no longer evident when compared to the control.

To determine whether oral infection with rCrPVs could also induce RNAi against target genes in *D. citri,* insects were fed on a solution containing 10^4^ TCID_50_ units/μL of each rCrPV. After 48 hours of feeding on the virus-containing solution, insects were transferred to a single citrus leaf system, where they were collected at different time points for gene expression analysis (RT-qPCR) and monitored for mortality (Figure 6A). Evaluation of viral loads during the course of infection showed that viral RNA levels decreased from 0 to 10 dpf, suggesting a partial clearance of the orally acquired recombinant virus, but viral RNA levels increased from 10 to 15 dpf, an indication of viral replication (Figure 6B). In alignment with these observations, and compared to rCrPV-GFP-fed insects, insects fed on artificial diets containing the rCrPV-IA-F and rCrPV-IA-R virions showed significant decreases (p < 0.05) in IA gene expression levels (36 and 30%, respectively) only at 15 dpf, the time point where viral RNA replication was detected (Figures 6C). From 0 to 10 dpf, no significant differences were observed in IA gene expression levels between the rCrPV-IA-F/R- and rCrPV-GFP-fed insects (Figure 6C). Additionally, no significant differences on the targeted gene expression levels between the rCrPV-IA-F- and rCrPV-IA-R-treated insects were observed in the feeding experiment. No significant differences in mortality rates were observed between rCrPV-IA-F/R- and rCrPV-GFP-fed insects during the monitored time frame (Figure 6D).

**Figure 6:**
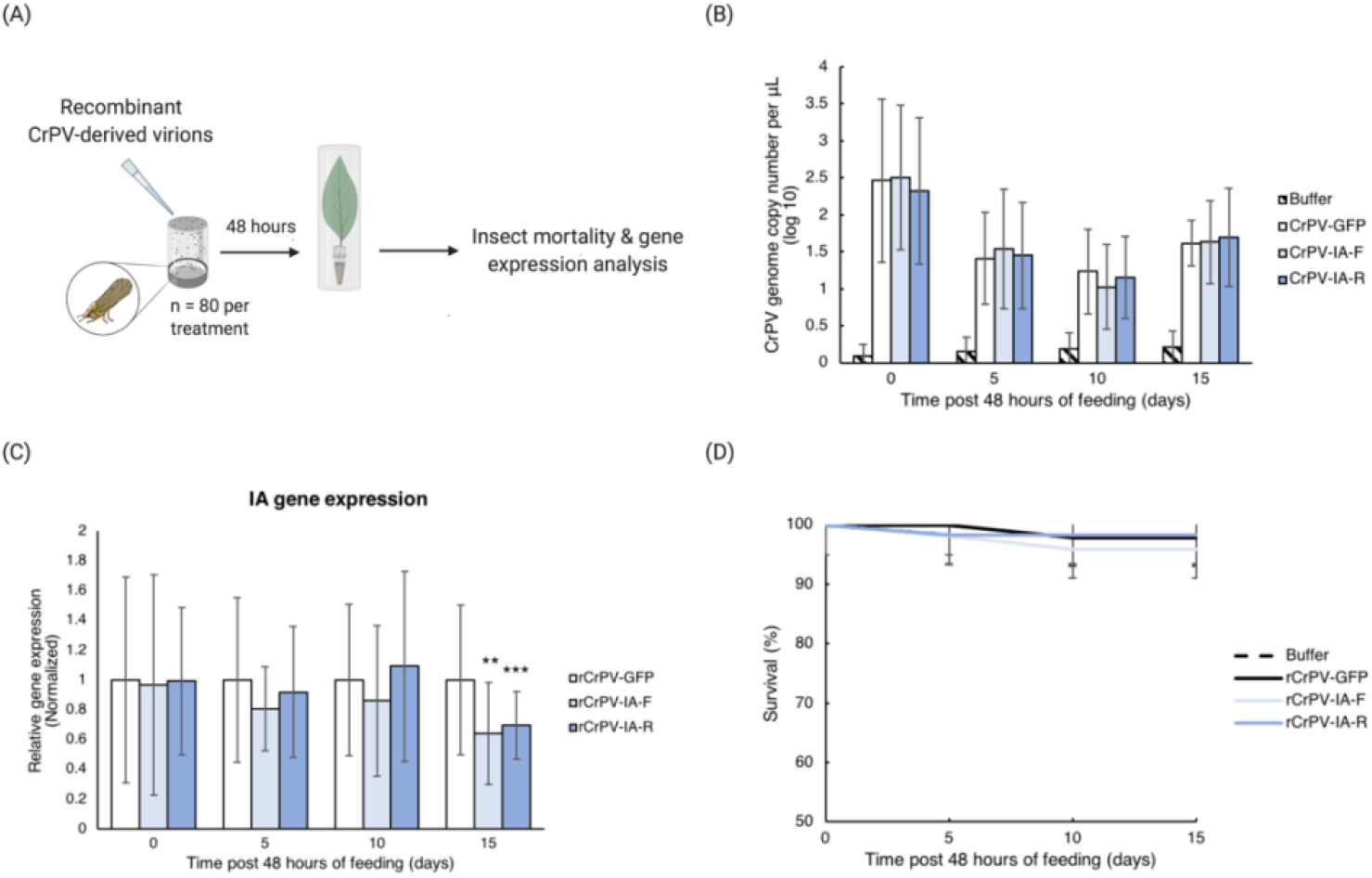
Virus-induced gene silencing (VIGS) using the recombinant Cricket paralysis virus (rCrPV) system for inducing RNA interference in *Diaphorina citri* by feeding delivery. (A) Schematic of the protocol. Adult insects (n = 80 total) were placed on an artificial diet feeding system consisting of rCrPV virions (10^4^ TCID_50_ units/μL) mixed in an artificial diet solution. Insects were allowed to feed for 48 hours and then transferred to a single citrus leaf system until further analysis. (B) Viral titers were checked at four time points after transfer of the insects from the virus-containing artificial diet solution to leaves and expressed as log10 genome copy number per μL (10 ng of RNA). Buffer-only fed insects were used as controls. Each time point represents data from ten individual insects; error bars indicate the standard error of the mean. (C) IA gene expression analysis in insects from (B) was performed by RT-qPCR. Insects fed on artificial diet containing rCrPV-GFP virions were used as controls and set as 1. IA gene-derived RNA levels were normalized against the *D. citri* actin gene. Bars represent the average IA gene RNA level of ten individual insects. Error bars indicate the standard error of the mean. ** = p < 0.005, *** = p < 0.0005 (two-tailed t-test). (D) Survival curve of adult insects fed on rCrPV containing artificial diet. Survival rates were monitored at four time points post feeding. Three biological replicates of n = 20 insects each per condition were analyzed. Error bars indicate standard error of the mean.

## Discussion

During the past two decades, RNAi-based technologies have been explored as tools for functional genomics studies but also for translational applications to help manage insect pests and/or plant pathogens. The majority of RNAi studies with insects have focused on using transformation-independent approaches, such as delivery of *in vitro* synthesized dsRNAs (the RNAi inducers) to the insect [4, 23, 25–29]. While these have yielded important fundamental information, some challenges of these approaches include: (1) the strategy employed for dsRNA delivery into the target insect, (2) the dsRNA dosage acquired by the insects, and (3) the dsRNA stability [13]. Additionally, most insects studied so far do not show a systemic dsRNA-mediated RNAi response, then if RNAi inducers are delivered to target insects via oral acquisition, RNAi effects may be limited to gut tissues [47]. In this work, we attempt to overcome some of these challenges and enhance opportunities for RNAi studies by using VIGS to induce an RNAi response in *D. citri*, an insect pest of critical agricultural importance.

Previously, Hajeri et al [48] used a plant virus, Citrus tristeza virus (CTV), as the backbone to engineer a VIGS vector that targets the abnormal wing disc (Awd) gene of *D. citri*. The authors demonstrated that insects fed on citrus plants infected with the recombinant CTV (carrying Awd-derived sequence) had lower expression levels of the Awd gene compared to controls, which resulted in malformed-wing phenotype in adult insects and higher insect mortality [48]. However, since most plant viruses do not replicate in insects (and this applies to CTV and *D. citri* as well), plant virus-induced gene silencing effects will depend on the insect feeding activity and on the levels of the respective interfering RNAs throughout the plant and acquired during insect feeding, and these effects will be likely limited to gut tissues [49].

The recent discovery of several *D. citri*-specific viruses through next generation sequencing [35] suggest the possibility to use VIGS to improve the efficiency, and likely systemic effects of RNAi-mediated gene silencing in *D. citri*, because these viruses spread naturally within the insect host [50]. However, due to a lack of established infectious clones for any known *D. citri*-specific virus, we used CrPV as a model virus to establish proof-of-concept for insect VIGS in *D. citri*. This is the first work showing CrPV infectivity and pathogenicity to *D. citri,* and shows that VIGS can be achieved, however the pathogenic effects induced by CrPV in *D. citri* are problematic. It is known that virus pathogenicity can be associated with many factors, including the ability of viruses to evade the host antiviral response, such as RNAi-mediated antiviral immunity [46]. CrPV, for example, encodes a potent silencing suppressor protein (1A), which likely acts as virulence determinant by interfering with the insect antiviral RNAi mechanism [46]. Since it is not known to what extent CrPV 1A silencing suppressor protein activity would affect RNAi studies in *D. citri*, we hypothesized that if the cleavage of the 1A protein was negatively affected during processing of the CrPV ORF1-encoded polyprotein, this might impact the ability of CrPV 1A to suppress the RNAi response in infected *D. citri*. Thus, based on Hahn and Palmenberg’s work [51], we performed mutational analyses on the CrPV DvExNPGP motif, which is the primary cleavage site for release of the 1A protein. The aa mutations introduced within the DvExNPGP motif either reduced or inhibited CrPV genome replication and resulted in a mutant (CrPV-DN*: DvExNPGP → NvExSPGP) that is attenuated compared to WT-CrPV, when infecting either *Drosophila* S2 cells or adult *D. citri* insects. In *D. citri*, this mutant had a lower replication rate, which probably contributed to a delay in insect mortality observed in CrPV-DN*-infected *D. citri* compared to the WT-CrPV-infected *D. citri*. Thus, we focused on using this mutant as the backbone for the construction of a recombinant CrPV-based VIGS system to specifically target *D. citri*.

To test the new CrPV-based VIGS tool, we selected a known *D. citri*-specific target: the inhibitor of apoptosis (IA) gene, which has been previously reported to induce early mortality in *D. citri* when targeted by a conventional RNAi approaches [4]. Insects either microinjected or fed with the new recombinant CrPV virions showed a decrease in the IA gene expression levels at time points coinciding with the initial detection of viral replication (at 5 hpi for injected insects, and at 15 dpf for fed insects), suggesting that the engineered CrPV-based VIGS system enables functional gene silencing in *D. citri*. However, our results also indicate that there might be several factors that need to be considered when engineering and applying the CrPV-based vectors for gene silencing in *D. citri*. The impact of the insert length on gene silencing, for example, may vary and needs to be investigated for each target. Galdeano et al. reported a decrease of 50% in the IA gene mRNA levels in *D. citri* fed on artificial diet containing an IA gene-derived dsRNA of 500 bp in length. In the present work, the silencing efficiency was reduced to 25-35% when *D. citri* were fed with 150 bp (maximum insert size tolerated by CrPV) *in vitro* synthesized dsRNAs of the same IA gene. On the other hand, a decrease of 50% in IA gene expression level was obtained when insects were injected with the same IA gene-derived dsRNAs (150 bp in length), suggesting that the dsRNA delivery method also affects the gene silencing efficiency in *D. citri*.

Another interesting observation was that the viral RNA accumulation in rCrPV-IA-microinjected insects were slightly higher compared with control insects (rCrPV-GF-microinjected insects), which could be related to the higher mortality also observed for these insects. However, there is no clear indication whether the higher viral titer and higher mortality rate are associated with effects of rCrPV-induced silencing on the target gene, since significant rCrPV-induced gene silencing was only observed during early stages of viral infection (5 hpi). These results highlight new questions involving VIGS and insect targets: How does target gene expression change in response to infection with a viral vector? Does RNAi-mediated silencing of a specific gene interfere (positively or negatively) with the replication of a VIGS viral vector in the insect? How does the interplay between target gene choice and viral replication affect the efficiency of a VIGS system in the insect? These questions need to be considered in future studies. Additionally, the method used for delivering the VIGS system is also important. Although CrPV can infect *D. citri* via oral feeding and, consequently, produces viral small interfering RNA [52], the viral replication rate is slower compared to when CrPV is delivered by microinjections. In this work, adult *D. citri* insects fed artificial diets containing the rCrPV-derived virions showed a significant decrease in the target gene expression levels only at 15 days post feeding, when an indication of virus replication was detected. Interestingly, analysis of viral RNA levels from the earlier time points suggested a partial clearance of the acquired recombinant virus. Similar results were also observed in a previous work, where authors reported clearance of orally acquired RNA viruses (including CrPV) in *Drosophila* flies [53]. Therefore, further investigations are needed to verify to what extent this virus clearance affects rCrPV-mediated gene silencing in *D. citri,* and also to explore how this tool can be improved.

Finally, we believe that the construction of a new insect virus-based VIGS system that is functional in inducing specific gene silencing in *D. citri* represents a valuable tool that can now be used as a model for future RNAi-based studies in this specific insect. The knowledge generated either in this study or in future studies using this developed system will help in understanding the interactions between insect virus-based VIGS systems and *D. citri,* and therefore will contribute to translating information to native *D. citri*-infecting viruses in order to explore new potential strategies for HLB control.

## Methods

### In vitro transcription and RNA transfection

CrPV virions were recovered from full-length CrPV cDNA cloned in pGEM vector driven by the T7 promoter (T7-pGEM-CrPV). Plasmids (1 ug) were linearized using BamHI restriction enzyme (NEB, Ipswich, MA, USA), column-purified using the DNA Clean & Concentrator −5 Kit (Zymo Research Corporation, Irvine, California, USA), and then used as template for *in vitro* transcription reaction using the T7 mMESSAGE mMACHINE Kit (Ambion, Austin, TX, USA), following the manufacturer’s instructions. Transcripts were phenol:chloroform-purified and the concentration and integrity were checked via nanodrop spectrophotometry and on a 1% denaturing glyoxal agarose gel.

*Drosophila* S2 cells were maintained in Schneider’s Insect medium (Sigma, St. Louis, MO, USA) supplemented with 10% fetal bovine serum (Sigma, St. Louis, MO, USA) and 5 U/ml of penicillin-streptomycin antibiotics (Gibco, Grand Island, NY, USA). Transfection of *in vitro*-synthesized RNAs into S2 cells was performed using TransMessenger reagent (Qiagen, Valencia, CA, USA) as per the manufacturer’s instructions. We used 1.6 ug of transcripts to transfect 3 × 10^6^ cells in a 12 well-plate. Transfected cells were incubated in 26 °C and monitored for cytopathic symptoms.

### Real-time quantitative RT-PCR (RT-qPCR) and Northern blot analysis

CrPV replication in S2 cells was checked by RT-qPCR and northern blot analysis. Transfected cells were resuspended in growth media and collected at time 0 (immediately after the final step of RNA transfection) and 4 days post transfection (when the presence of the cytopathic symptoms was clearly observed). Cells and supernatant were separated by low-speed centrifugation (4.000 rpm) for 10 min at 4 °C. Total RNAs were isolated from the pellet and supernatant using TRIzol reagent (Life Technologies, Carlsbad, CA, USA). RNAs were reverse transcribed using the High-Capacity cDNA Reverse Transcription Kit (Applied Biosystems, Carlsbad, CA, USA), following the manufacturer’s protocol. qPCR was performed to detect the CP gene of CrPV using 1X of SsoAdvanced Universal SYBR Green Supermix (Bio-Rad, Foster City, CA, USA), 300 nM of each specific primer (CrPV-CPF/CrPV-CPR, Table S1), 1 μL of the 1:5 diluted cDNA and water up to 10 μL total volume. The thermocycling conditions were the following: 95 °C for 3 min, followed by 40 cycles of 95 °C for 10 s and 58 °C for 30 s. The same procedure was done with a dilution series of CrPV plasmid in order to establish the standard curve. The plasmid copy number was calculated as described by Plumet and Gerlier [54]. The standard curve was then used to estimate the CrPV genome copy number in the tested samples.

For northern blot analysis, 1 ug of total RNA isolated from the pellet and supernatant was denatured with glyoxal at 55 °C for 30 min, electrophoresed on a 1% agarose gel and transferred to the Hybond-NX membrane (GE Amersham, Piscataway, NJ, USA) by capillary transfer. RNAs were fixed to the membrane by cross-linking in UV light and blots were stained with methylene blue. The membranes were hybridized with ^32^P-labeled probes to detect the positive-strand RNA. For making the probes, amplified and gel-purified DNA fragments containing the T7 promoter and covering nt 8284-8672 (CP) of the CrPV genomic RNA were used as templates for *in vitro* transcription reaction using T7 RNA polymerase (Ambion MAXIscript T7 Transcription Kit, Vilnius, Lithuania) in the presence of [α-32P] UTP, according to the manufacturer’s instructions. After hybridization, the membranes were washed once in 2x SSC (1× SSC is 0.15 M NaCl plus 0.015 M sodium citrate) and 0.1% SDS at RT for 15 min, once in 0.5x SSC/0.1% SDS at RT for 15 min and once with 0.1x SSC/0.1% SDS at 65 °C for 15 min. The membranes were exposed to Premium X-Ray film (Phenix Research Products, Candler, NC, USA) for hybridization detection.

### Virion purification: SDS-PAGE, TEM and viral titration

For virion purification, virions recovered from transfected S2 cells were propagated by inoculating 1 mL of infected S2 cells culture in 50 mL culture flasks containing 3 × 10^6^ of cells in 6 mL of same culture media, followed by incubation at 26 °C until detection of the cytopathic symptoms. The entire cell cultures were then harvested and submitted to a 10 min centrifugation at 4.000 rpm and 4 °C. The supernatants were transferred to new tubes and centrifuged for 30 min at 8.000 rpm and 4 °C. Virus particles in the supernatant were pelleted by ultracentrifugation at 45.000 rpm for 45 min (Beckman 70.1 Ti Rotor) over a pad of 15% sucrose in 10 mM Tris-HCl buffer (pH 7.4). Pellets were resuspended in same buffer and overlaid on CsCl gradient (from bottom to top: 1.6, 1.5, 1.4, 1.3 and 1.2 g/cm^3^ of CsCl in 10 mM Tris-HCl buffer) for 4 hrs, at 50.000 rpm and 11 °C (Beckman SW65 Ti Rotor). The virion band was extracted using an 18‐ gauge needle syringe, diluted in 9 mL of 10 mM Tris-HCl buffer and centrifuged for 45 min at 45.000 rpm (Beckman 70.1 Ti Rotor). Pellets were suspended in the same buffer and analyzed by SDS-PAGE using 12.5% polyacrylamide gel, followed by a Coomassie Brilliant Blue R-250 (Fisher Biotech) staining (Sambrook and Russell David, 1989). An aliquot of the purified virions (5 μL) was loaded onto Formvar-carbon coated grids, left for 2 min and drained with filter paper. The grids were stained using 5 drops of 2% uranyl acetate, then drained and air-dried. The stained grids were examined with a JEOL 2100F transmission electron microscope at 200 kV accelerating voltage.

Virus titer was measured by the end-point dilution method. Purified virions were filtered in a 0.22 μm syringe filter and used in 10-fold serial dilutions. S2 cells (10^4^ cells per well in a 24-well plate) were inoculated with the viral dilution series (four replicates each dilution) and the cytopathic effects were analyzed 48 hours post inoculation. The viral titer was calculated as TCID_50_ according to a published method [54].

### Viral infection in D. citri

Virus-free *D. citri* colonies were maintained on young healthy citrus (*Citrus macrophylla*) plants in cages housed in a climate-controlled greenhouse at 25 °C. The infection experiments were conducted using approximately 20 day-old insects (n = 60) by intrathoracically injecting approximately 300 nL of a 10^3^ TCID_50_ units/μL virus suspension in 10mM Tris buffer, pH 7.4. Same volume of 10 mM Tris buffer (no virus) was also injected as a mock-infected control. Microinjected insects were kept in a single leaf system and monitored daily for live/dead insect counts and symptoms detection. Microinjected insects were collected at time 0, 3 and 6 dpi for RT-qPCR and northern blot analysis. For each time point, 3 live insects were pooled in three biological replicates for total RNA extraction. RT-qPCR and northern blot analyses were performed as described above.

### Construction of CrPV mutants: in vitro and in vivo analyses

The T7-pGEM-CrPV plasmid was used as template to generate independent mutants (Figure 2B) by using two partially complementary overlapping mutagenic primers (Table S1). Reverse PCRs were performed in two rounds using CloneAmp HiFi PCR premix (Clontech, Mountain View, CA, USA). The first round was performed adding the forward and reverse primers in two independent reactions and consisted of 3 cycles of 98 °C for 10 seconds, the respective primers annealing temperature for 15 seconds and 72 °C for 1 min. The reactions with the forward and reverse primers were mixed and then subjected to a second round of 16 cycles of amplification following the same conditions. The amplicons were gel purified and 100 ng of the mutated linearized plasmids were re-ligated by using the NEBuilder HiFi DNA Assembly Cloning Kit (NEB, Ipswich, MA, USA), following the manufacturers’ protocol, and transformed into *E. coli* (DH5α). Plasmids were purified from the transformed colonies using the QIAprep Spin Miniprep Kit (Qiagen, Valentia, CA, USA) and the mutated regions were confirmed by Sanger sequencing.

Mutated plasmids were digested with BamHI restriction enzyme (NEB, Ipswich, MA, USA), purified and used for *in vitro* transcription reaction as described above. Transfected cells were monitored daily for cytopathic symptoms and collected at time 0 and 96 hours post transfection (hpt). The viral RNA accumulation of the CrPV mutants from time 0 to 96 hpt were compared with the WT-CrPV by RT-qPCR and northern blot analysis, following previously described procedures.

Virions from a selected CrPV mutant (CrPV-*DN) that showed lower RNA accumulation compared with the WT-CrPV were purified from S2 cells cultures as described. Viral titer was calculated by end-point dilution method and same amounts of WT and CrPV mutant (approximately 300 nL of a 10^3^ TCID_50_ units/μL virus suspension in 10mM Tris buffer, pH 7.4) were intrathoracically microinjected in *D. citri*. Sixty insects approximately 20-days-old were used for each treatment. Microinjected insects were kept in a single leaf system and monitored daily for mortality. Viral RNA accumulation in microinjected insects was checked by RT-qPCR at 0, 3 and 6 dpi exactly as described above.

### Recombinant CrPV-based VIGS in D. citri

The attenuated mutant CrPV-DN* from this work, was used as template for construction of the recombinant CrPVs. The insertion site of foreign sequences in the CrPV-DN* vector was created downstream of the CrPV DvExNPGP motif (modified to NvExSPGP in the CrPV-DN* mutant). A duplicated DvExNPGP motif was inserted downstream of the insertion site. Different nucleotide lengths of GFP gene-derived sequences (57, 102, 150 and 201 nt in length) were amplified from a previously constructed GFP expression vector and individually cloned in the insertion site of the CrPV-based vector, following previously described procedures. The stability of the new recombinant CrPV vectors carrying different insert lengths was checked by recovering the recombinant virions in S2 cells, as already described. Among the tested insert lengths that did not affect the virus stability, the largest one (150 nt) was selected as the optimum insert length for the CrPV-based VIGS system. To verify whether these short sequences can still trigger RNAi responses in *D. citri*, fragments of 150 nt were amplified from the GFP gene (control) and from the selected *D. citri* target gene (IA, GenBank accession number: XM_008469919) using primers containing the sequence of the T7 promoter at the 5’ ends. PCR products containing the T7 promoter at the 5’ ends of each strand were used for dsRNA synthesis following the MEGAscript RNAi kit (Ambion, Foster City, CA USA) protocol. The *in vitro* synthesized dsRNAs were confirmed on 1% agarose gel and delivered to *D. citri* insects either by microinjection (200 ng per insect) or oral feeding (300 ng/μL of dsRNA solution). Microinjected insects (n = 60) were left in a single citrus leaf assay for approximately 17 hours and fed insects (n = 60) were left on artificial diets containing the respective dsRNAs for 5 days. Ten insects from each experiment were collected and used for single insect RNA extraction. Gene expression analysis (to validate dsRNA-mediated gene silencing) was performed as described in the following paragraphs.

The fragments of 150 nt in length were amplified considering both forward (rCrPV-IA-F) and reverse (rCrPV-IA-R) orientations of the *D. citri* IA gene (they do not represent the exact same region of the IA gene because the reverse-oriented sequence had to be adjusted to not interfere with the ORFs of CrPV) and cloned in the insertion site of the CrPV-based vector. The recombinant CrPV carrying a fragment of 150 nt derived from the GFP gene (non-*D. citri* target gene) was used as control in this study. The recombinant CrPV virions were recovered in S2 cells and checked by SDS-PAGE as described. To verify whether the inserts were maintained stably in the virion RNA, we analyzed the insertion site region by RT-PCR followed by Sanger sequencing.

For the recombinant CrPV-based VIGS experiment, adult *D. citri* insects (approximately 20 days-old; n = 80 for each treatment) were infected with the respective rCrPVs virions (rCrPV-GFP; rCrPV-IA-F and rCrPV-IA-R) via both feeding and intrathoracically microinjection. A mock experiment containing only buffer was performed as negative control in both assays. Microinjected insects (approximately 300 nL of 10^4^ TCID_50_ units/μL per insect) were kept in a single citrus system and 10 insects were collected at time 5, 17 and 40 hpi for target gene expression analysis. Feeding assays were performed following an artificial diet feeding system described in Galdeano et al. [4]. Insects were fed to rCrPV virions suspension (10^4^ TCID_50_ units/μL) mixed in an artificial diet solution [10% sucrose (w:v); 0.1% green and 0.4% yellow food dyes (McCORMICK & CO)] for 48 hrs and then kept in single citrus leaf assay. Ten fed insects were collected at time 0, 5, 10 and 15 days post 48 hrs of feeding for target gene expression analysis. Both microinjected and fed insects were also daily monitored for mortality.

Collected insects were submitted to single insect RNA extractions using TRIzol reagent (Life Technologies, Carlsbad, CA, USA), treated with RQ1 RNase-free DNase I (Promega, Madison, WI, USA), and purified by phenol:chloroform extraction. cDNAs were generated using the High-Capacity cDNA Reverse Transcription Kit (Applied Biosystems, Carlsbad, CA, USA), following the manufacturer’s protocols, and qPCR was performed in three technical repeats as described above using primers to detect the *D. citri* IA gene (Table S1), designed to bind outside the targeted region. Relative expression levels of the target *D. citri* gene were normalized to the endogenous *D. citri* actin gene and calculated using Pfaffl’s method (Pfaffl 2001). The statistical difference between the test and the control groups was calculated by *t-test* (p < 0.05). RT-qPCR analysis was also performed to detect the CrPV CP gene and calculate the expression CrPV RNA genome copy number in both microinjected and fed insects. Results are representatives of two independent experiments.

## Acknowledgements

This work was financially supported by grants from the US Department of Agriculture (grants 13-002NU-781; 2015-70016-23011, AP19PPQS&T00C234) and the University of California. We wish to thank our laboratory colleagues from UC Davis and from the Contained Research Facility (CRF), where the described experiments were performed. We would like to especially thank Dr. S. W. Ding (UC Riverside) for have kindly provided the CrPV plasmid; and Dr. Fei Guo (UC Davis BioEM facility) for helping with the electron microscopy. There are no conflicts of interest to disclose regarding this work.

---

**Figure S1:**
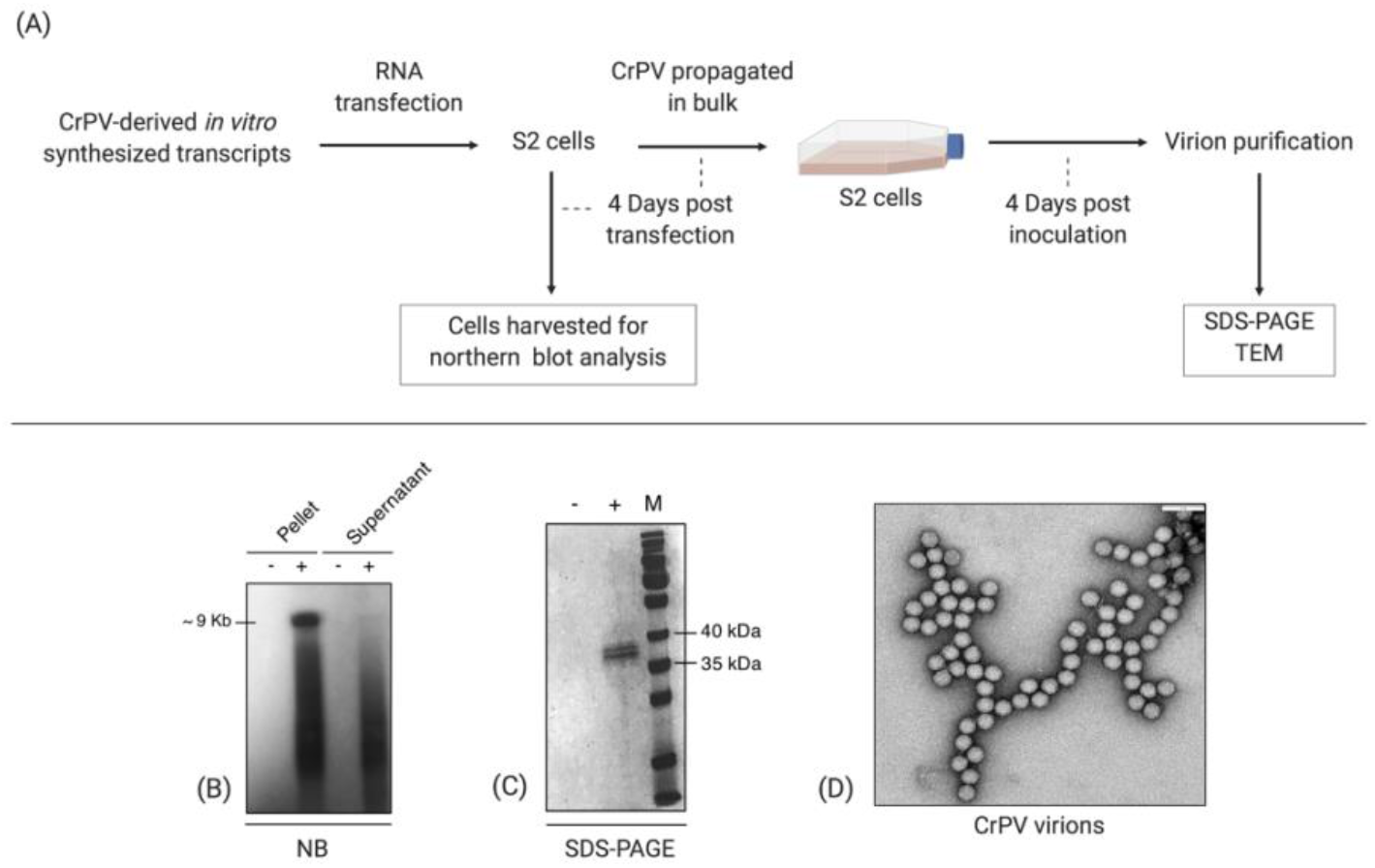
Recovery of Cricket paralysis virus (CrPV) virions in *Drosophila melanogaster* Schneider line 2 (S2) cells. (A) CrPV virions were propagated in S2 cells by transfecting *in vitro*-transcribed RNAs derived from a CrPV infectious clone. Viral replication was confirmed by northern blot (NB) analysis on the total RNA extracted from cultures of transfected S2 cells (from pelleted cells and supernatant separately) (B). CrPV virion preparations were verified using SDS-PAGE (C) and transmission electron microscopy (TEM) (D). (−) Mock, reagent-only transfected cells; (+) CrPV-transfected cells; M, Page ruler prestained protein ladder.

**Figure S2:**
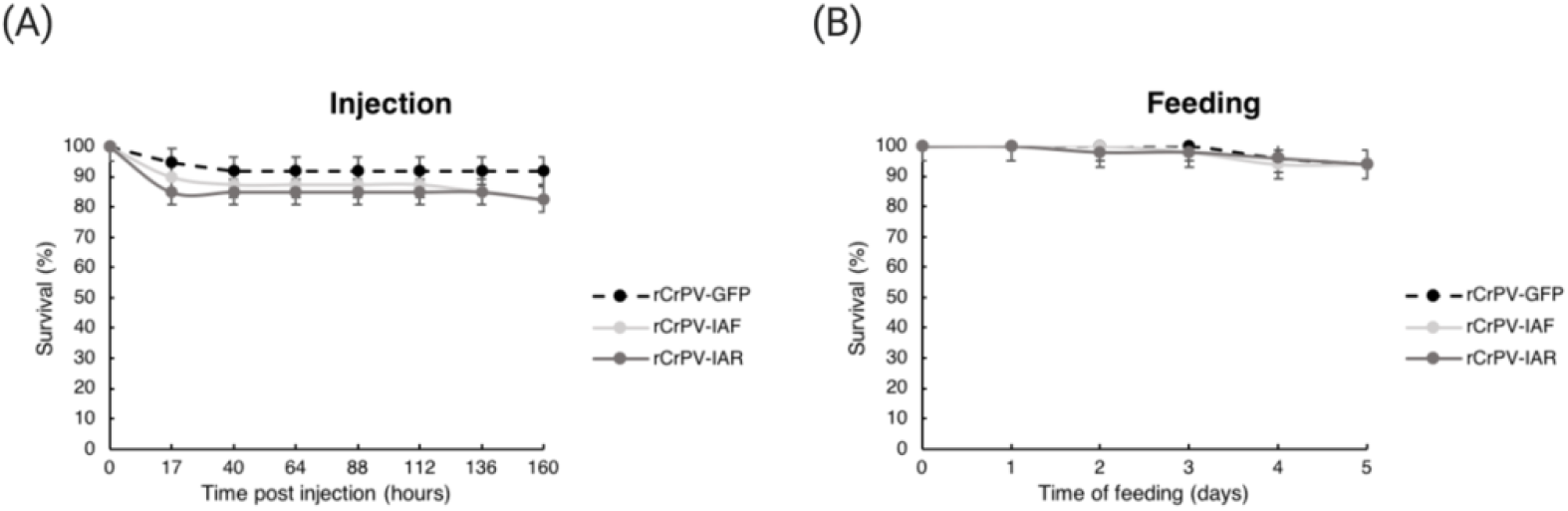
Impact of inhibitor of apoptosis (IA) gene silencing on survival rates of insects (*Diaphorina citri*) treated with IA-derived dsRNAs. (A) Survival curve of adult insects injected with IA dsRNAs. Survival rates were monitored at eight time points post injection. Three biological replicates of n = 20 insects each per condition were analyzed. Error bars indicate standard error of the mean. (B) Survival curve of adult insects over five days of feeding on artificial diet containing IA dsRNAs. Survival rates were daily monitored during feeding. Three biological replicates of n = 20 insects each per condition were analyzed. Error bars indicate standard error of the mean.

**Figure S3:**
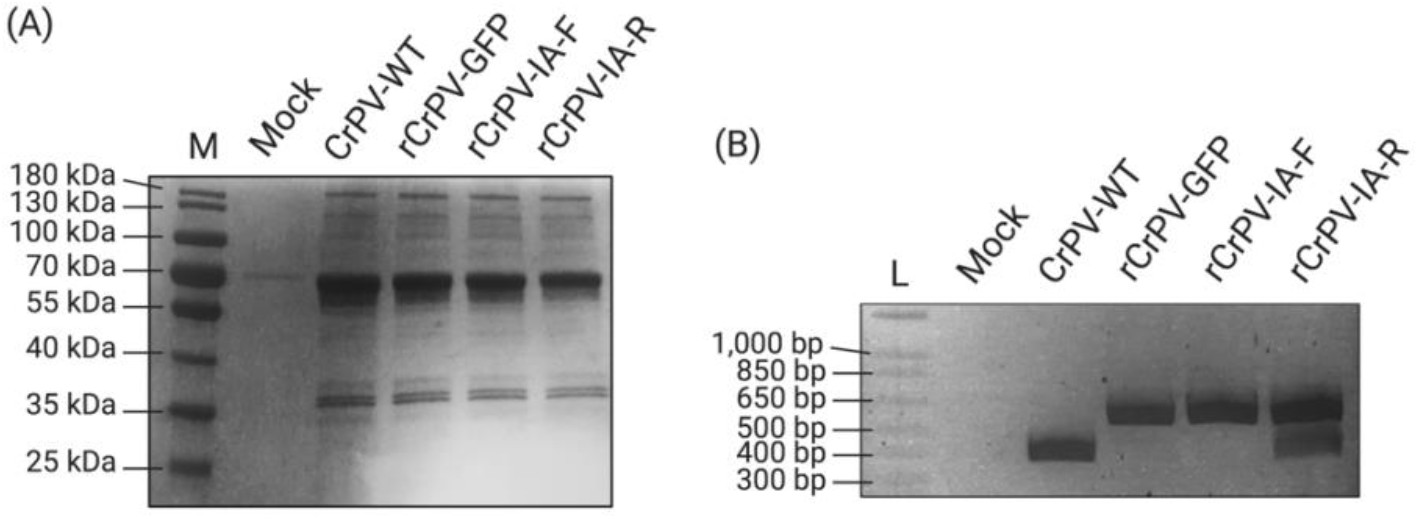
Recombinant CrPV (rCrPV) virions carrying the target gene-derived sequences were propagated in *Drosophila* S2 cells and checked by SDS-PAGE (A) and by RT-PCR of the insertion site region to confirm the retention of the target-derived inserts (B). M, Page ruler prestained protein ladder; L, 1 Kb plus DNA ladder.

**Table S1:**
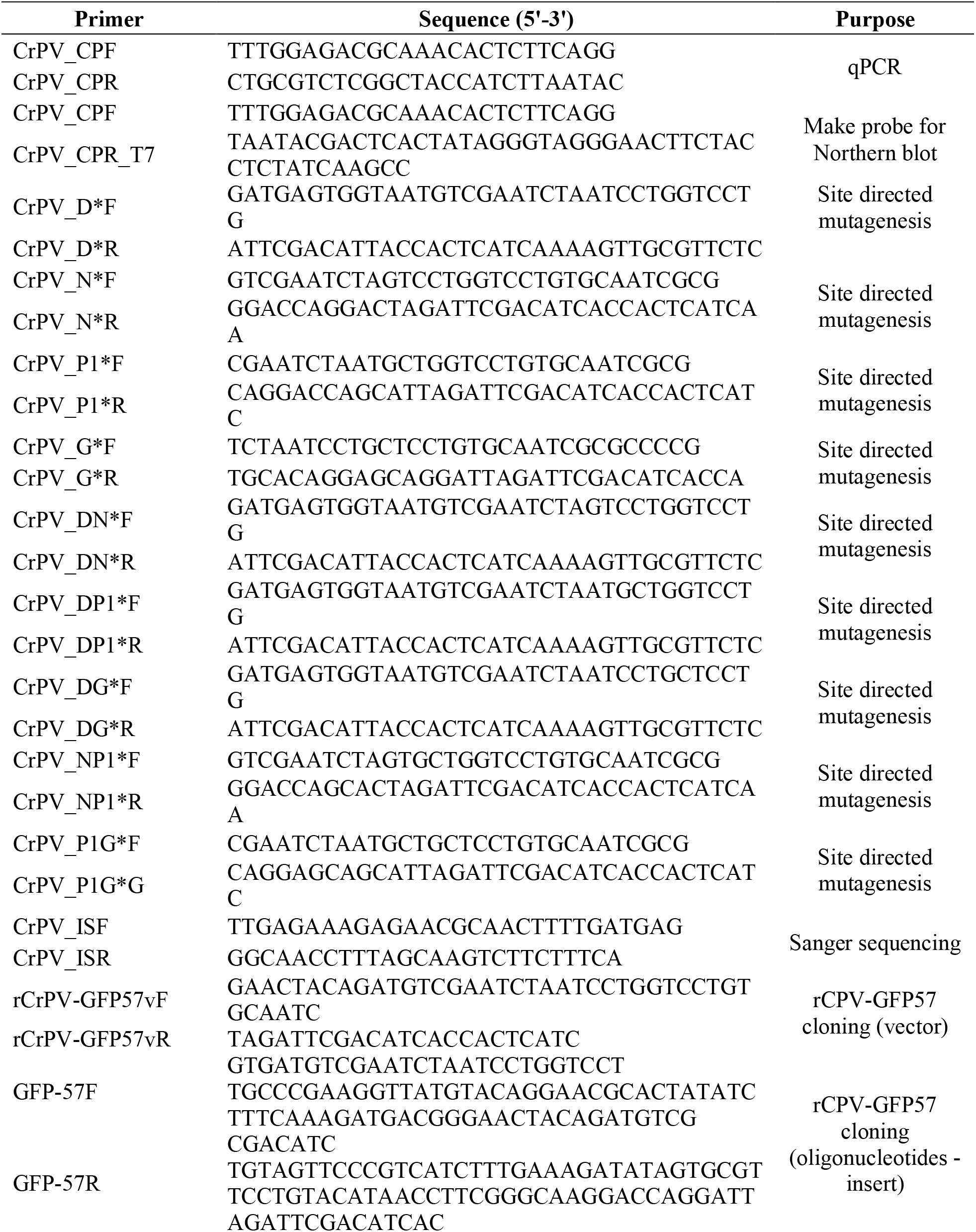
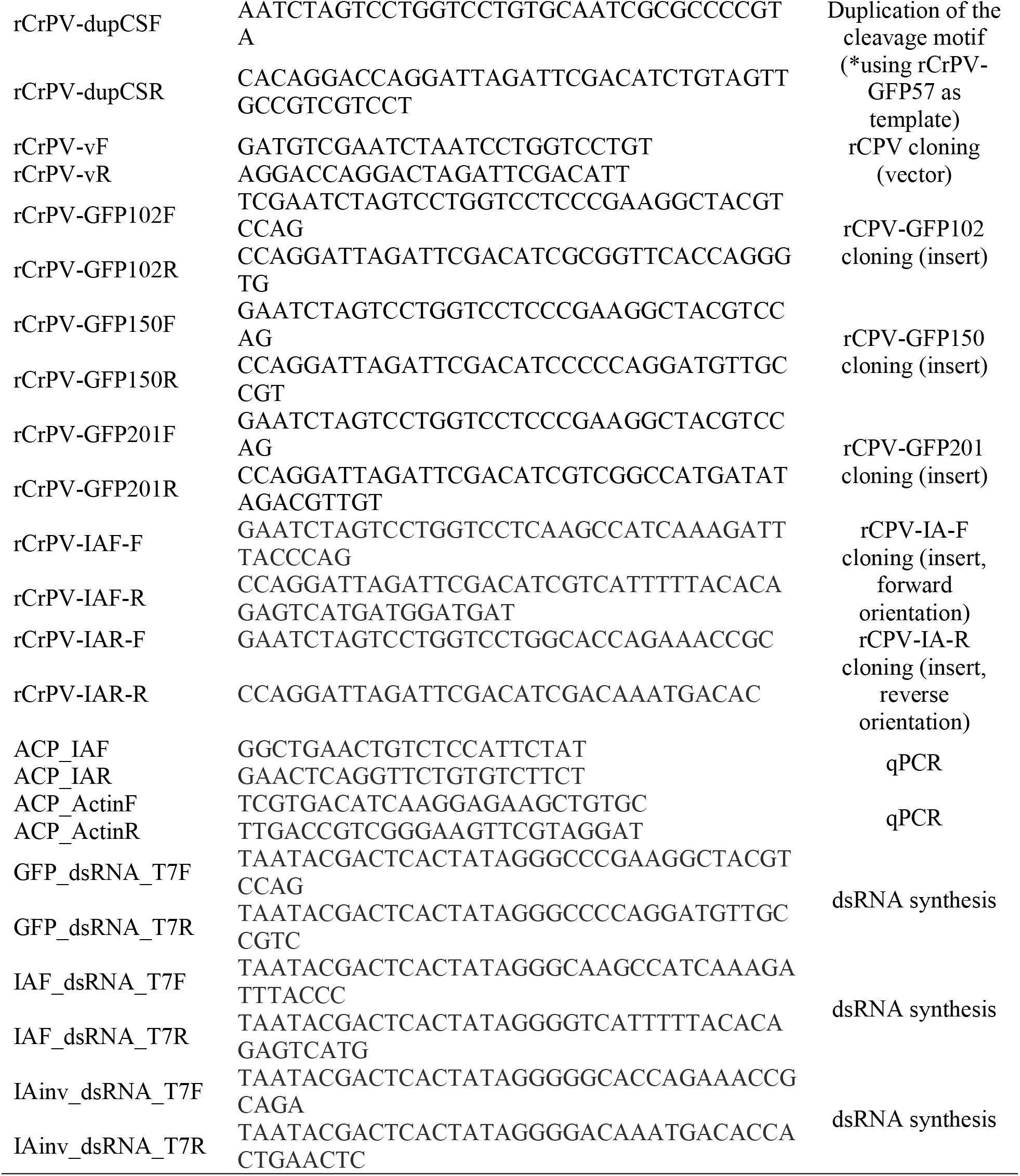
List of primers used in this study. The purpose of each primer is presented.

## Notes

### Competing Interest Statement

The authors have declared no competing interest.

